# Potent broad-spectrum anti-coronaviral frameshift inhibitors from virtual screen of RNA binding

**DOI:** 10.1101/2025.09.17.676934

**Authors:** Krishna Neupane, Rohith Vedhthaanth Sekar, Sandaru M. Ileperuma, Eileen Reklow, Sneha Munshi, Sara Ibrahim Omar, Sahar Arbabimoghadam, Jennifer A. Peterson, Jack A. Tuszynski, Tom C. Hobman, Michael T. Woodside

## Abstract

Coronavirus genomes contain an RNA pseudoknot that directs −1 programmed ribosomal frame-shifting (−1 PRF) to control expression of viral proteins crucial for replication. Ligands that inhibit −1 PRF can thus attenuate viral propagation and have potential as drugs for limiting coronavirus infections. To search for novel small-molecule frameshift inhibitors with anticoronaviral activity, we computationally screened over 14 million compounds for binding to the SARS-CoV-2 pseudoknot, followed by experimental validation of the top hits for inhibition of −1 PRF and viral replication. We identified multiple potent −1 PRF inhibitors, effective at nM concentrations, some of which significantly suppressed SARS-CoV-2 replication in cell culture. Several compounds also inhibited −1 PRF in multiple representative bat coronaviruses, indicating broad-spectrum activity. These results showcase the promise of viral RNA structures like frameshift-stimulatory pseudoknots as targets for broad-spectrum antiviral drugs.

## INTRODUCTION

When it emerged in 2019, Severe Acute Respiratory Syndrome coronavirus 2 (SARS-CoV-2) caused the most disruptive pandemic in the last century, with world-wide excess mortality estimated at ∼15 million in the first two years of the pandemic.^1^ The rapid development of effective vaccines helped to lower mortality, and several drug treatments (including antibody therapies, nucleotide-analog polymerase inhibitors, and protease inhibitors) exist to treat infections.^2^ However, novel variants of the virus with increased ability to evade neutralizing antibodies from vaccines, prior infections, or therapeutics continue to emerge,^3–5^ and the evolution of resistance to existing drugs threatens to reduce their effectiveness.^6,7^ There thus remains an important need to develop novel drugs for treating COVID-19 infection, especially ones that can be used in combination therapies to minimize the likelihood of drug resistance. Furthermore, novel coronaviruses (CoVs) can be expected to pose an ongoing threat for future zoonotic outbreaks, owing to increased human contact with reservoirs such as bats,^8^ underlining the need for drugs with activity against a broad spectrum of CoVs.

All existing approved antivirals target proteins in the virus or host. However, elements within the viral RNA itself offer potential targets that can be exploited for novel antiviral therapeutics. One such target is the RNA pseudoknot stimulating −1 programmed ribosomal frameshifting (−1 PRF), a process essential for replication of all CoVs.^9^ In −1 PRF (Fig. 1A), the ribosome is directed to shift upstream by 1 nucleotide (nt), into the −1 reading frame, by a tripartite signal in the mRNA consisting of a 7-nt slippery sequence at which the frameshift occurs, a ∼5–7-nt spacer, and a structure in the mRNA (typically a pseudoknot, as in CoVs) that stimulates the frameshift.^10,11^ In CoVs, −1 PRF is required to translate the open reading frame encoding proteins needed for viral RNA transcription and replication, including the viral polymerase.^9^ Modulating −1 PRF levels can significantly reduce CoV replication,^12–14^ making −1 PRF an attractive therapeutic target. Moreover, the frameshift signal is among the most conserved regions of CoV genomes,^15^ thus reducing the likelihood of mutations conferring drug resistance.^16^ In addition, targeting −1 PRF is orthogonal and complementary to existing strategies of targeting proteins such as the viral polymerase or proteases, offering the potential for effective combination therapies. Finally, many CoV pseudoknots share structural similarities that could allow for binding of ligands with activity against −1 PRF in a broad spectrum of CoVs.^17,18^

**Figure 1:**
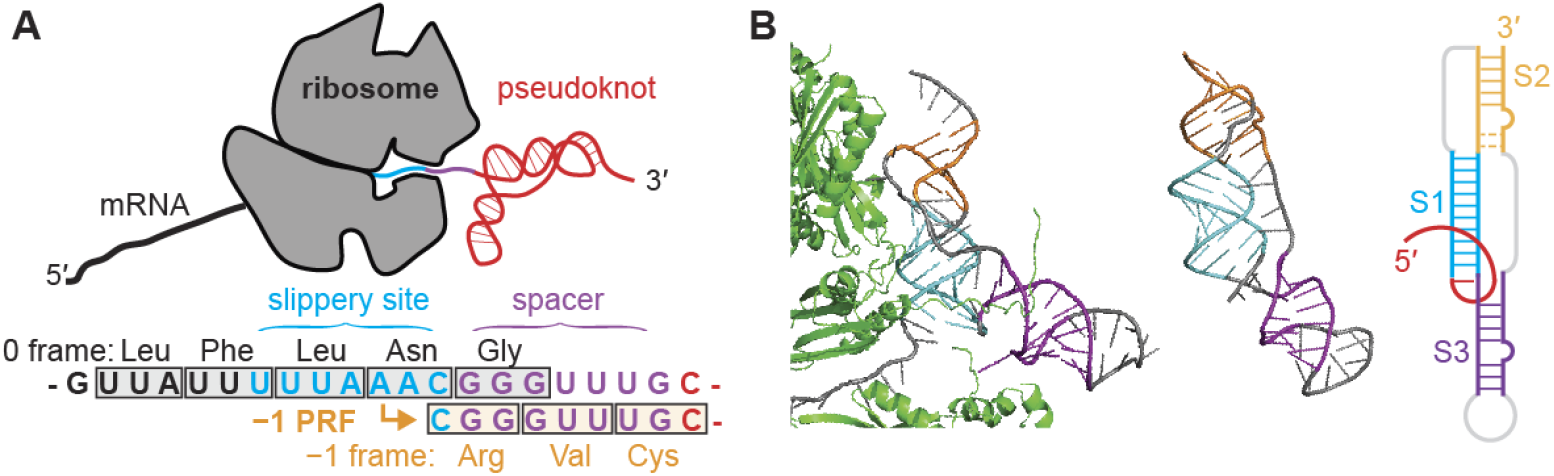
−1 Programmed ribosomal frameshifting. (A) Schematic of −1 PRF. −1 PRF is stimulated by a structure in the mRNA (for SARS-CoV-2, a 3-stem pseudoknot, red) downstream of a slippery sequence (cyan). The shift is reading frame changes the viral polypeptide sequence (bottom inset). (B) Structural models of SARS-CoV-2 frameshift-stimulatory pseudoknot (right: secondary structure; left: structure of pseudoknot arrested on ribosome (ref. 20); center: computational model of structure (ref. 23).

Previous work has identified −1 PRF in SARS-CoV-2 as a potential drug target. SARS-CoV-2 has been confirmed to undergo −1 PRF,^19^ and suppression of −1 PRF by small-molecule drugs has been shown to inhibit viral infectivity.^14^ Most importantly, extensive studies have revealed structural and dynamical properties of the frameshift-stimulatory pseudoknot relevant for structure-based drug discovery.^20–27^ The pseudoknot was found to contain 3 stems, rather than 2 as is much more common, and to feature an unusual topology in which the 5′ end is threaded through a ring formed by the 3-helix junction when the pseudoknot interactions create stem 2 (Fig. 1B). The complexity of the 3-stem junction region creates a number of potential binding pockets with the potential to support both high-affinity and high-specificity interactions.^28^ Furthermore, the rarity of the 5′-threaded topology—reported previously only in a few viral non-coding RNAs^29,30^—suggests that ligands specific to the SARS-CoV-2 pseudoknot may have low propensity to interact with off-target RNAs.

Small-molecule ligands modulating −1 PRF in SARS-CoV-2 and other human CoVs have been reported in several studies,^14,16,17,19,31–35^ as have anti-sense oligomers specific to CoV pseudoknots.^27,36,37^ Small-molecule ligands were in most cases found through empirical screens of relatively small chemical libraries (∼10^3^–10^4^ compounds), using translation of a reporter mRNA construct to identify −1 PRF modulators. Notable results from such empirical screens include merafloxacin, an antibacterial fluoroquinolone reported to reduce −1 PRF in SARS-CoV-2 by up to ∼90% with IC_50_ (concentration yielding half-maximal inhibition) of ∼ 20 μM;^14^ nafamostat, a serine protease inhibitor reported to reduce −1 PRF in SARS-CoV-2 by ∼50–70% with IC_50_ ∼ 0.5 μM;^17^ and KCB261770, a novel compound reported to reduce −1 PRF in SARS-CoV-2 by ∼70% with IC_50_ ∼ 0.5 μM.^31^ Intriguingly, all of these compounds were found to be active against frameshift signals from more than one CoV, suggesting possible broad-spectrum anti-CoV activity. In addition, a virtual screen of a ∼100,000-entry library for docking against a computational model of the SARS-CoV-1 pseudoknot^34^ found a compound with IC_50_ ∼ 0.8 μM that was later confirmed to inhibit −1 PRF in SARS-CoV-2.^16,19^ However, so far, no inhibitors have been reported that are effective in the low-nM concentration range needed for practical application as therapeutics.

To find more potent −1 PRF inhibitors, we used experimental and computational models of the SARS-CoV-2 pseudoknot structure^20,23^ to screen a library of ∼14 million small-molecule compounds *in silico* via blind docking and molecular dynamics simulations. Experimental tests revealed multiple hits that inhibited −1 PRF *in vitro* in SARS-CoV-2. Several of these hits had high potency, with IC_50_ ∼ 50 nM or below, and some were also effective at suppressing viral replication in a cell culture model of infection. Furthermore, several hits showed strong potential for broad-spectrum anti-CoV activity by inhibiting −1 PRF caused by frameshift signals representative of the sequences found in a wide range of CoVs from bats, a key zoonotic reservoir of CoVs. These results, from one of the largest computational screens yet reported on any RNA target, identify potent new frameshift inhibitors with promising potential as anti-CoV drug candidates with broad-spectrum activity.

## RESULTS

To leverage the particular advantages of frameshift-stimulatory pseudoknots as drug targets in CoVs, we focused on searching for pseudoknot-binding ligands. The SARS-CoV-2 pseudoknot exhibits conformational heterogeneity, forming two distinct fold topologies.^22^ Given that increased conformational heterogeneity is linked to higher −1 PRF levels,^38,39^ potent frameshift inhibitors should stabilize the most commonly occupied structure instead of less-occupied conformers. In the case of SARS-CoV-2, the dominant conformer was the 5′-threaded one, as seen in both static structural studies of the pseudoknot^20,27^ and measurements of its folding dynamics.^22^ We therefore chose to use models of the threaded conformer for docking calculations to find pseudoknotbinding −1 PRF inhibitors (Fig. 1A). Specifically, we chose two models: (1) a cryo-EM model of the pseudoknot solved in complex with the ribosome, where the ribosome was arrested on the slippery sequence during translation in cell extract,^20^ as the most biologically relevant static structure resolved so far; and (2) a computational model of the pseudoknot in isolation^23^ that is in good agreement with measurements of single-molecule folding dynamics.^22^ Both models are in reasonable agreement with the solution structure of the pseudoknot assayed by small-angle x-ray scattering.^40^

We screened the ZINC15 library^41^ computationally (Fig. 2A), looking at ∼14 million drug-like compounds (after excluding those that were listed as not commercially available or that did not have a three-dimensional representation), to sample a large chemical space. Because the binding sites of candidate ligands were unknown, we divided the pseudoknot into 5 overlapping potential binding volumes covering most of the surface area of the two pseudoknot models (Fig. 2B). We then performed blind docking using rDock,^42^ an algorithm that has previously been calibrated for RNA ligands,^43,44^ calculating the docking score for each ligand against all 5 docking sites on both structural models. Ligands were then ranked by their top docking score (among all 5 sites on each model), and the top-scoring hits were tested in 200-ns long all-atom molecular dynamics simulations to verify the persistence of the binding in the predicted pose (rejecting compounds that did not remail bound). A sample binding pose after molecular dynamics simulations is shown in Fig 2C for one of the ligands.

**Figure 2:**
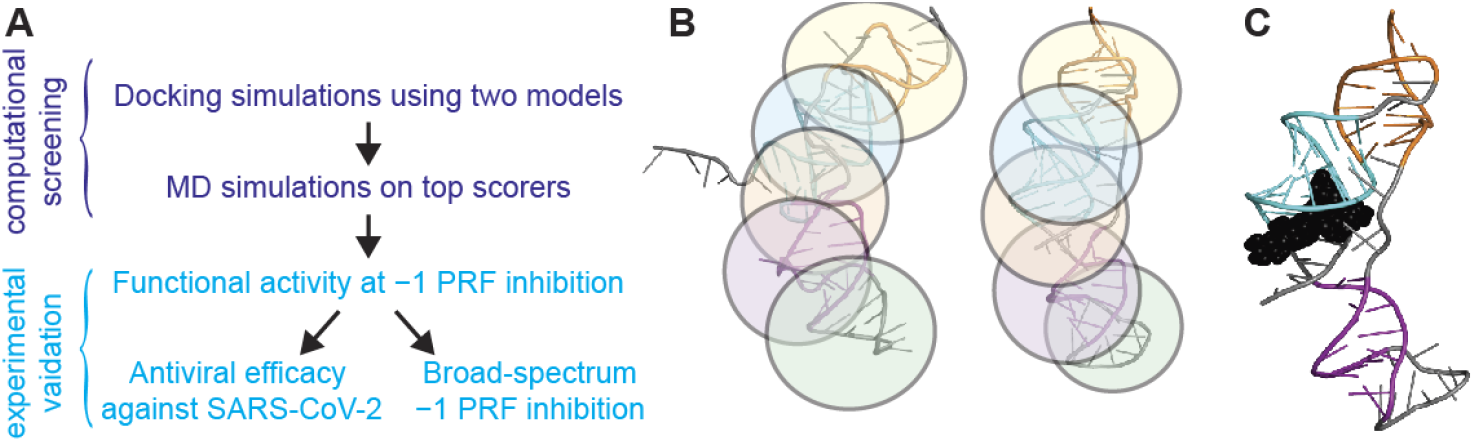
Virtual screening for binding to SARS-CoV-2 pseudoknot. (A) Flowchart of screening process. (B) Two structural models (left: from cryo-EM of pseudoknot arrested on ribosome; right: from computational modeling) were used for blind docking, with each structure divided into 5 binding regions for the calculations. (C) Sample binding pose of a ligand on the pseudoknot (representative structure from most occupied cluster in molecular dynamics simulation).

For experimental validation of the virtual screening hits, we chose the top 131 compounds that we could purchase commercially, and measured the effect of each compound on −1 PRF using a cell-free dual-luciferase assay as in previous work.^16,17,19^ A reporter mRNA consisting of the *Renilla* luciferase gene in the 0 frame upstream of the CoV frameshift signal followed by the firefly luciferase gene in the −1 frame was translated in cell lysate, and the −1 PRF efficiency was obtained from the ratio of luminescence emitted by the two enzymes, as compared to controls with 100% and 0% firefly luciferase read-through. The inhibition level was then quantified by comparing the result in the presence of 20 µM compound to that without compound. The results show that the hits from the virtual screening are heavily enriched in frameshift inhibitors, as desired (Fig. 3A, Supporting Data S1): over half of the compounds (68 of 131) inhibited −1 PRF at the 2-σ level (Fig. 3A, red). A substantial fraction of the compounds, just under 1 in 4, generated a moderate effect size (∼30– 50% reduction of −1 PRF), and ∼7% caused strong inhibition (∼50% reduction or more). In contrast, screens of compound libraries that have not undergone any kind of selection for interactions with the machinery of −1 PRF have found that very few compounds show any effect on −1 PRF,^14,17^ indicating the effectiveness of the virtual screening process at selecting compounds that preferentially modulate −1 PRF.

**Figure 3:**
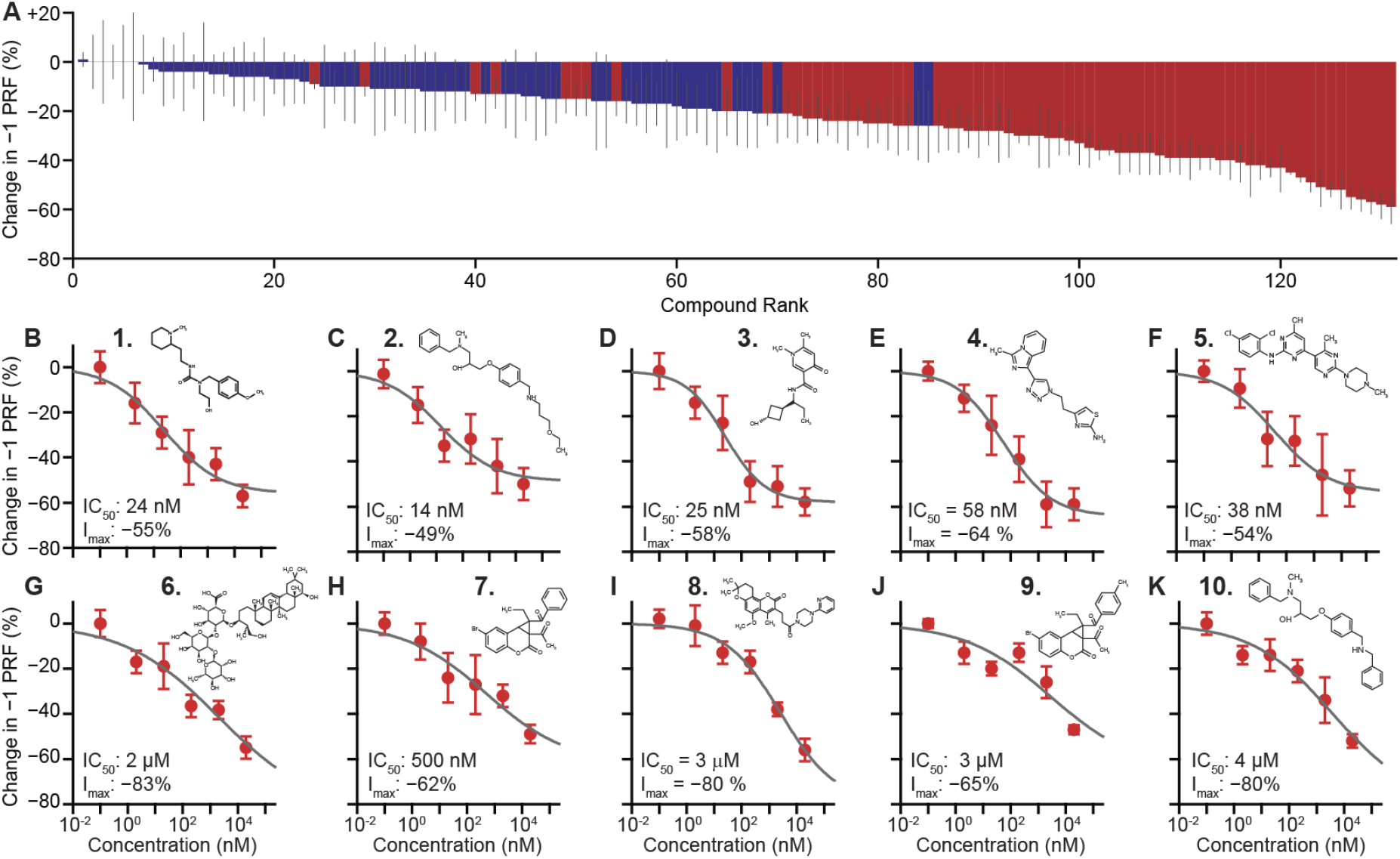
Evaluating screening hits for ability to inhibit−1 PRF. (A) Survey of the change in −1 PRF stimulated by the SARS-CoV-2 frameshift signal that was induced by 20 µM of each compound, as measured by a dual-luciferase assay. Results are shown ranked by increasing inhibitor strength. Compounds reducing −1 PRF by more than 2σ are shown in red. (B–K) Dose-response of the 10 most effective inhibitors from (A). Insets: compound structures. Error bars represent s.e.m. from 3–11 replicates.

For the top ten strongest inhibitors, we next characterized their dose-dependent effect by repeating the dual-luciferase assays at different ligand concentrations between 0.1 nM and 20 µM (Fig. 3B–K). We fit the dose-dependent −1 PRF inhibition for each compound to determine the inhibition potency (IC_50_) and maximum inhibitory effect size (I_max_). We found that I_max_ values were in the range of roughly −50 to −80%. Five of the compounds had IC_50_ below 60 nM, with three of them in the 15–25 nM range, 10–1000-fold lower than for the most effective inhibitors reported to date, whereas for the others IC_50_ was in the range 0.5–4 µM, comparable to several previously reported inhibitors.

To assess the ability of the strongest inhibitors to attenuate viral infections, we infected 293T Ace2 cells^45^ with SARS-CoV-2, using multiplicity of infection (MOI) of 0.5, after treating them with varying concentrations of −1 PRF inhibitor (0.1–10 µM). Treated cells were compared to negative controls (no inhibitor) and cells treated with 1 µM remdesivir, a nucleotide-analog polymerase inhibitor approved for treatment of COVID-19,^46^ as a positive control. Two of the compounds (numbers 2 and 6) significantly reduced viral titers in the cell media (Fig. 4A) as well as the amount of viral RNA in cell lysates (Fig 4B), in a dose-dependent manner. For compound 2 (Fig. 4, blue), a noticeable reduction in viral RNA was seen at 0.1 µM but infectious titers were only slightly lower, within the uncertainty; at 1 µM, viral RNA was reduced ∼40-fold and infectious titers ∼20-fold. For compound 6 (Fig 4, brown), a ∼3–4-fold reduction in viral RNA at 0.1 and 1 µM was accompanied by a similar decrease in infectious titers, and 40–50-fold decreases in both were seen at 10 µM. Neither compound was cytotoxic at the maximum dose used in the antiviral activity assays (Fig. 4C).

**Figure 4:**
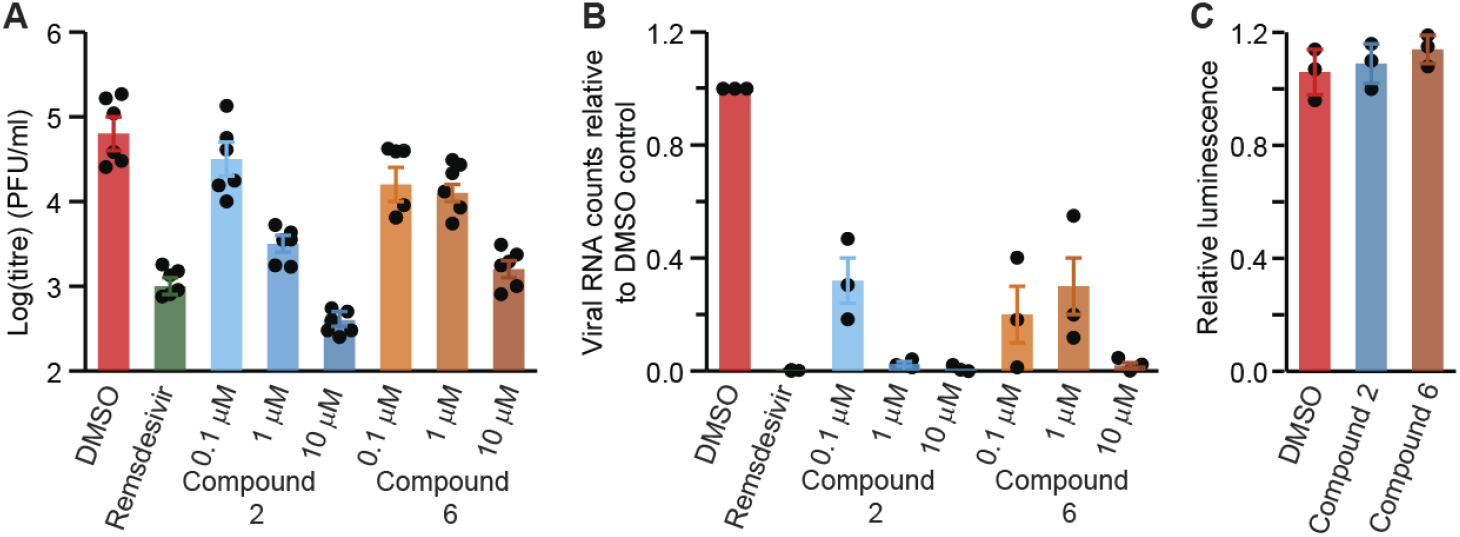
Effect of −1 PRF inhibitors on SARS-CoV-2 replication. 293T Ace2 cells treated with inhibitors or DMSO alone were infected with SARS-CoV-2, then media and total RNA were harvested for plaque assay and qRT-PCR to measure viral titers and relative levels of viral genomic RNA, respectively. (A) Compounds 2 and 6 reduced SARS-CoV-2 titers in a dose-dependent way compared to DMSO negative control, as measured by plaque assay. Remdesivir (1 µM) was used as a positive control. (B) Compounds 2 and 6 reduced the amount of viral RNA relative to the DMSO control (defined as 1.0) in a dose-dependent way as assayed by reverse transcriptase qPCR. Remdesivir (1 µM) was used as a positive control. (C) Cell viability assay using luminescent measure of ATP concentration revealed no cytotoxic effects for 10 µM compounds 2 and 6, as compared to the DMSO negative control. Error bars represent s.e.m. from 3–6 replicates.

Finally, we explored the ability of these compounds to inhibit −1 PRF stimulated by a broad range of CoV frameshift signals, to test whether the compounds had broad-spectrum activity against other CoVs (Fig. 5). We used a panel of bat-CoV frameshift signals, since bats have been the ultimate source of most human CoVs. Previous work found that the high conservation of the frameshift signal in CoVs results in a small number of characteristic structures for bat-CoV pseudoknots.^18^ We chose frameshift signals from two bat CoVs having pseudoknots representative of the structures found in bat beta-CoVs (GLGC2 and Vs-CoV-1; a third cluster of bat bet-CoVs is SARS-like and hence already represented by SARS-CoV-2), and two having pseudoknots representative of structures found in bat alpha-CoVs (Anlong-44 and Neixiang-64). We also included measurements using the frameshift signal from HIV-1, to test for the specificity of the −1 PRF inhibitors; only compounds 9 and 7 showed a change in −1 PRF statistically different from 0 for HIV-1, and even then the effects were relatively small (∼15–20%). All compounds except compound 2 were effective to at least a moderate extent (over 30% inhibition) against at least one of the bat-CoV frameshift signals; several were so against two, and compound 6 against all four, indicating a very broad spectrum of activity at inhibiting −1 PRF in CoVs.

**Figure 5:**
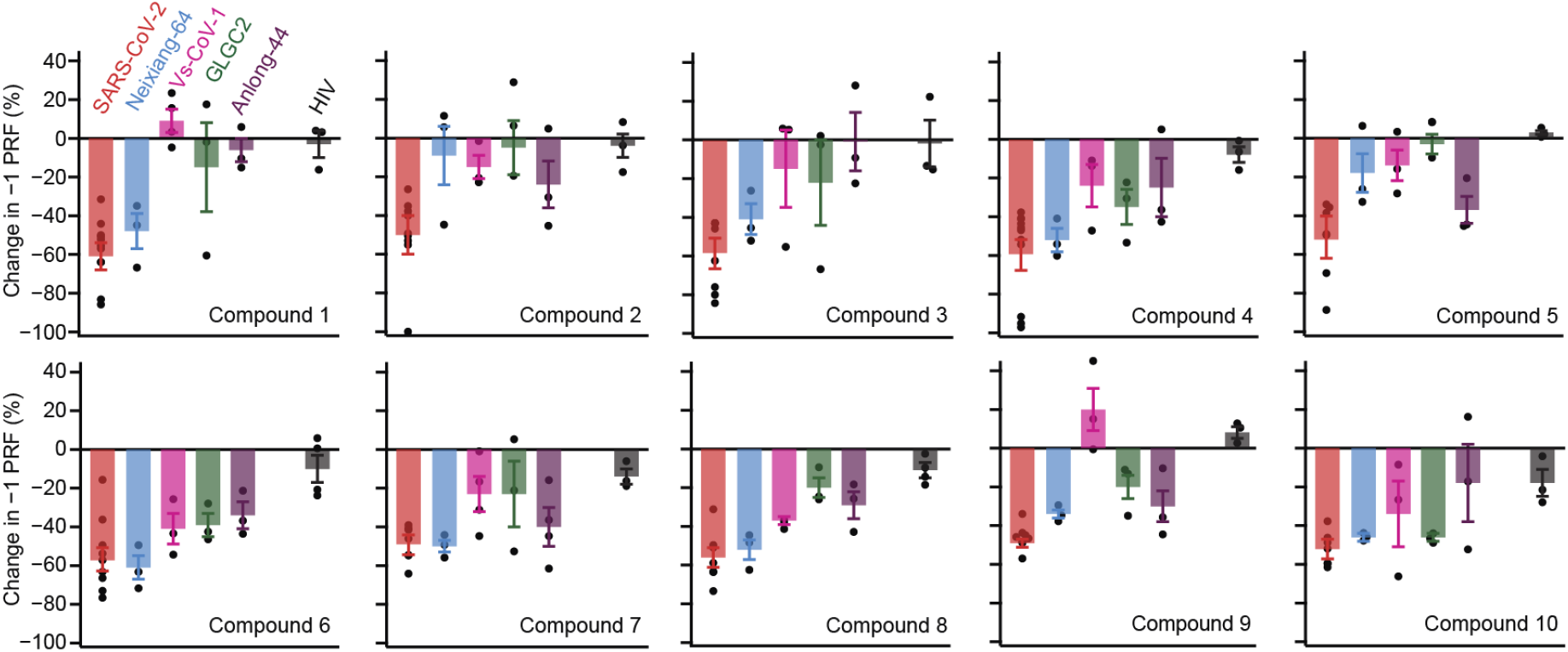
Testing activity against a broad-spectrum of bat-CoV frameshift signals. Each of the 10 compounds studied in Fig. 3B–K was tested for activity against 4 frameshifting signals representative of the range of sequences found in bat CoVs. The frameshift signal of HIV was included as a control to test specificity. Error bars represent s.e.m. from 3–11 replicates.

## DISCUSSION

Targeting viral RNA structures is a very promising area for developing novel antivirals, as they represent a diverse pool of potential therapeutic targets, with structures distinct from host RNAs and complex enough to generate binding pockets with high affinity and specificity. However, viral RNAs remain under-explored as antiviral targets. Pseudoknots that regulate translational frameshifting in viral RNA are among the few RNA structures that have been targeted with small-molecule drugs, but efforts to date have yielded only a handful of compounds that modulate −1 PRF significantly in SARS-CoV-2, none of them potent at the nM-scale concentrations needed for practical therapeutics. By searching computationally through a much larger chemical library than explored previously, we discovered multiple compounds that are not only effective at reducing −1 PRF by two-fold or more—a level that should be sufficient to dramatically reduce viral replication^13,14^—but also potent at much lower concentrations than the strongest inhibitors reported previously for SARS-CoV-2: 10 to 1000-fold lower, in the ∼10–50 nM range. The proportion of hits from the virtual screen that are strong −1 PRF inhibitors was also orders of magnitude higher than for previous empirical high-throughput screens (∼7% vs ∼0.02–0.1%), suggesting that the computational screening step is very effective at identifying active inhibitors. Notably, none of the compounds enhanced −1 PRF, unlike as found in empirical screens,^14,17^ consistent with the hypothesis that targeting the dominant threaded conformer of the pseudoknot should reduce −1 PRF by decreasing conformational heterogeneity.^38,39^

These results also broaden the structural diversity of the small molecules verified to act as −1 PRF inhibitors for SARS-CoV-2. One notable chemical difference between previously reported inhibitors and those discovered here is that whereas the former consist largely of polyaromatic, flat compounds (Fig. S1), the latter have more three-dimensional structures owing to the presence of sp^3^ centers. Given that 4 of the 5 most potent new inhibitors contain one or more sp^3^ centers, we speculate that this feature may be important for achieving high potency. Several of the new inhibitors also feature oxygen-substituted rings that make them more electron-rich. Other novel features not seen in previously reported inhibitors include cyclopropane rings (compounds 7 and 9), a cyclobutane ring (compound 3), and a steroid-sugar (compound 6, the natural product soyasaponin I). To explore the structural diversity more systematically, we used molecular fingerprinting to cluster the set of 131 inhibitors we tested experimentally (Fig. S2). We found that most compounds were chemically distinct from all others (including the inhibitors found previously), but a few were part of clusters of chemically similar compounds (including five of the compounds in the top ten: compounds 2, 5, 7, 9, and 10 from Fig. 3), indicating that the virtual screening consistently scored these scaffolds quite highly.

The high potency of −1 PRF inhibition in the cell-free assay was only partly reflected by the potency in the cell-culture infection model. For compound 2, at 100 nM the viral RNA counts were reduced by ∼3-fold and the infectious titers were reduced by ∼2-fold (within the errors), whereas from previous data on infectious titers as a function of −1 PRF inhibition^14^ the results from Fig. 3C suggest viral titers would be expected to be reduced by ∼6-to 8-fold at this concentration. The discrepancy was much smaller for compound 6, where infectious titers were reduced by ∼4-fold at 100 nM, close to the expected 4-to 6-fold reduction. Given that −1 PRF inhibition and infectious viral titers were seen to correspond tightly for SARS-CoV-2 in previous work,^14^ this discrepancy in potency suggests that the effective inhibitor concentration inside the cells may be lower than the treatment concentrations, possibly due to membrane permeability effects or removal by efflux pumps. Chemical modification of the compounds to increase membrane permeability and/or evade pumping by P-glycoprotein or other pumps may thus increase the antiviral potency.

Turning to mechanism of action, the inhibitors are expected to act by binding to the frameshift-stimulatory pseudoknot, given that the screen used to discover them was based on computational docking to the pseudoknot. Using microscale thermophoresis to look for binding to purified SARS-CoV-2 pseudoknot did indeed show that compound 2 binds the pseudoknot, with dissociation constant *K*_D_ of ∼3 μM (Fig. S3). However, the signal-to-noise ratio was relatively low, raising the uncertainty in *K*_D_. For compound 6, MST binding signals were even smaller, such that affinity could not be quantified. These results suggest that binding to the pseudoknot on its own may be weaker than expected from the IC_50_ values, given *K*_D_ estimated at more than two orders of magnitude higher than IC_50_ for compound 2, and that the changes in diffusion upon compound binding are sufficiently small that reliable quantification of affinity is difficult. We speculate that the discrepancy between *K*_D_ and IC_50_ for compound 2 may arise because the compound inhibits −1 PRF with high potency by interacting not just with the pseudoknot but also synergistically with other factors like the ribosome, and that its affinity for the pseudoknot alone is thus weaker than would be expected. Such a situation is plausible given putative binding sites near the pseudoknot-ribosome interface. A similar discrepancy between *K*_D_ and IC_50_ was also seen previously for an inhibitor of −1 PRF in SARS-CoV-1.^34,47^

Finally, we note that despite docking only against the frameshift-stimulatory pseudoknot from SARS-CoV-2 in the computational screen, almost all the top inhibitors were active against at least one of the frameshift signals representative of those found in bat CoVs. Eight of the ten were at least moderately effective against Neixiang-64 (an alpha-CoV), five against Anlong-44 (another alpha-CoV), and three each against GLGC2 and Vs-CoV-1 (both beta-CoVs). Remarkably, one compound (compound 6) was effective against all four of the bat CoVs tested. Intriguingly, two of the compounds sharing a similar chemical scaffold (compounds 2 and 10) showed a different spectrum of activity: compound 2 was the most potent against −1 PRF for SARS-CoV-2 but inhibited −1 PRF for only one of the bat-CoVs, whereas compound 6 was less potent against −1 PRF for SARS-CoV-2 but had more broad-spectrum activity, inhibiting −1 PRF for 3 of the bat CoVs. These results highlight the potential for medicinal chemistry derivatization of the inhibitors to improve their spectrum of activity. Notably, however, all inhibitors were generally inactive against the HIV-1 frameshift signal (where the stimulatory structure is a hairpin^48,49^), implying that the inhibitors were likely acting through the frameshift-stimulatory pseudoknot. The ability of so many frameshift inhibitors for SARS-CoV-2 to inhibit frameshifting in other CoVs—despite not including those CoV pseudoknots in the computational docking screen—presumably arises from the fact that the CoV pseudoknots share certain structural features, such as the similar stem 1 and threaded 5′ end structures noted in previous work.^18^ This unplanned versatility of SARS-CoV-2 frameshift inhibitors highlights the potential for broad-spectrum anti-CoV activity of −1 PRF inhibitors.

## CONCLUSIONS

High-throughput computational screening of ∼14 million small-molecule compounds followed by experimental testing of the top hits discovered multiple compounds that inhibited −1 PRF *in vitro* in SARS-CoV-2. Several inhibitors were highly potent, with IC_50_ in the nM range; some were also effective at suppressing viral replication in a cell culture model of infection. Moreover, several compounds inhibited −1 PRF *in vitro* for multiple pseudoknots representative of the variety found in bat CoVs, the zoonotic source of most human CoVs. These results, from one of the largest computational screens yet reported on any RNA target, identify potent new frameshift inhibitors with promising potential as anti-CoV drug candidates with broad-spectrum activity.

## METHODS

### Virtual screening

We used the ZINC 15 library^41^ as the source of ligands for virtual screening. We first filtered the library for all compounds having information on their three-dimensional representation and listed as “in-stock” under the Purchasibility filter, resulting in ∼14 million compounds. The resulting library of compounds was loaded into Molecular Operating Environment (version 2019.01) and the database wash function used to protonate the compounds and generate tautomers (protonation menu set to “enumerate” with maximum limit set to 5 and minimum concentration 20%) in preparation for docking. Docking simulations were performed with rDock,^42^ using two receptor models: a structural model of the SARS-CoV-2 pseudoknot solved by cryo-EM from a ribosome-mRNA complex in which the ribosome was stalled at the slippery site in the frameshift signal,^20^ and a computational model of the pseudoknot in isolation.^23^ As the most effective binding sites for frameshift inhibition were unknown, we chose to dock each compound across the entire pseudoknot, dividing each of the two structural models into 5 docking zones with radius of cavity mapping region set to 15 Å with centers for each site located roughly equidistant from each other as shown in Fig. 2B. For each zone on each structural model, we generated 200 docking runs for every entry in the prepared database of ligands. For each receptor model, compounds were ranked based on the score of their highest-scoring pose across all five zones, and the top-scoring compounds were taken for the next round of analysis.

To test the stability of the ligands in their predicted binding poses for each receptor, we ran all-atom explicit water molecular dynamics simulations of the ligand-pseudoknot complex for 200 ns using Amber18.^50^ We used the Antechamber program within AmberTools to parameterize the ligands, and the ff99OL3 force field^51^ to parameterize the RNA pseudoknot. The system was solvated using optimal point charge water boxes, with minimum distance between the complex and the box boundaries set to 12 Å. The solvated systems were first neutralized using sodium ions, then their salinities were adjusted to 0.15M NaCl using Joung-Cheatham monovalent ion parameters.^52^ The solvated systems were subjected to energy-minimization and then heated to 310 K with restraints of 10 kcal/mol/Å^2^ on the ligand-pseudoknot system, before gradually removing the restraints and subsequently simulating the unrestrained system for 200 ns at constant pressure. Ligands that dissociated from the pseudoknot within the 200-ns trajectory (defined as having root mean square deviation greater than 2.5 Å in the last 50 ns of the trajectory) were rejected as false positives.

### Preparation of mRNA constructs

(1) Constructs for dual-luciferase assays of −1 PRF were prepared as described previously.^16,19^ Briefly, we used a dual-luciferase reporting system based on a plasmid containing the sequence for *Renilla* luciferase and the multiple cloning site (MCS) from the plasmid pMLuc-1 (Novagen) upstream of the firefly luciferase sequence in the plasmid pISO (addgene). The frameshift signals for SARS-CoV-2, bat CoVs, and the specificity control (HIV-1) were cloned into the MCS between the restriction sites PstI and SpeI. Three different construct types were made for each frameshift signal. First, we made a construct for assaying −1 PRF efficiency, containing the frameshift signal with slippery sequence (UUUAAAC) and pseudoknot in the 0 frame and upstream of the firefly luciferase gene in the −1 frame, so that expression of the latter was dependent on −1 PRF. We also made two controls: (1) a negative control to measure the background firefly luciferase luminescence (0% firefly luciferase read-through), in which the slippery sequence was mutated to include a stop codon (UUGAAAC); and (2) a positive control to measure 100% firefly luciferase read-through, in which the slippery sequence was disrupted (UAGAAAC) and the firefly luciferase gene was shifted to the 0 frame. Sequences of frameshift signals for all constructs are listed in Table S1. We amplified transcription templates from these plasmids by PCR, using a forward primer that included the T7 polymerase sequence as a 5′ extension to the primer sequence.^16,19^ We transcribed mRNAs for dual-luciferase measurements from these templates *in-vitro* using the MEGAscript T7 transcription kit (Invitrogen), and purified them with the MEGAclear transcription clean-up kit (Invitrogen). All mRNAs were polyadenylated (including 30 A’s at the 3′ end) before translation but left uncapped.

### Dual-luciferase assays of −1 PRF

The −1 PRF efficiency was measured using a cell-free dual-luciferase assay.^53^ Briefly, for each construct, 1.2 µg of mRNA transcript was heated to 65 °C for 3 min and then incubated on ice for 2 min. The mRNA was added to a solution mixture containing amino acids (10 µM Leu and Met, 20 µM all other amino acids), 17.5 µL of nuclease-treated rabbit reticulocyte lysate (RRL, Promega), 5 U RNase inhibitor (Invitrogen), and brought up to a reaction volume of 25 µL with water. The reaction mixture was incubated for 90 min at 30 °C. Luciferase luminescence was then measured using a microplate reader (Turner Biosystems). First, 20 µL of the reaction mixture was mixed with 100 µL of Dual-Glo Luciferase reagent (Promega) before reading firefly luminescence, then 100 µL of Dual-Glo Stop and Glo reagent (Promega) was added to the mixture to quench firefly luminescence before reading *Renilla* luminescence. The −1 PRF efficiency was calculated from the ratio of firefly to *Renilla* luminescence, *F*:*R*, after first subtracting the background *F*:*R* measured from the negative control (which defines the signal expected for 0% −1 PRF) and then normalizing by *F*:*R* from the positive control (which defines the signal expected for 100% −1 PRF).

To quantify the effects of the inhibitors on −1 PRF efficiency, compounds were added to the reaction volume at the desired final concentration for each construct (frameshift signals from SARS-CoV-2, bat CoVs, and HIV, as well as all positive and negative controls). Compounds were dissolved in DMSO, leading to a final DMSO concentration in the assays of 1% by volume. The results were averaged from 3– 11 replicates, as described previously.^16,19^ The *Renilla* and firefly luciferase luminescence levels were similar for the different frameshift signals. The compounds did not affect the luminescence levels for *Renilla* luciferase, indicating that they did not interfere with translation, but rather their effects were specific to −1 PRF. IC_50_ and I_max_ were found by fitting the concentration-dependent inhibition, *I*(*x*), to *I*(*x*) = I_max_(1 – [1 + (*x*/IC_50_)^*d*^]^−1^), where *d* parametrizes the sharpness of the inhibition curve.^54^

### Cell culture, infection, and antiviral assays

293T Ace2 cells^45^ were cultured in Dulbecco’s modified Eagle’s medium (DMEM) supplemented with 100 U/mL penicillin and streptomycin, 1 mM 4-(2-hydroxyethyl)-1-piperazineethanesulfonic acid, 2 mM glutamine, and 10% heat-inactivated fetal bovine serum (FBS) at 37 °C in 5% CO_2_. To test antiviral activities of compounds, 293T Ace2 cells were plated in 96-well plate (3.0 × 10^4^ cells per well) and incubated 24 hrs at 37 °C. Compounds dissolved in DMSO were mixed in DMEM mix (1% final DMSO concentration) and then added to cells at the desired final concentration (0.1–10 µM) for 1 hour before infection with SARS-CoV-2 (CANADA/ON-VIDO-01/2020 isolate) using a multiplicity of infection (MOI) of 0.5. After 24 hr, cell media were harvested for plaque assays and lysed with Machery-Nagel RA1 buffer for quantitative RT-PCR analyses, respectively. SARS-CoV-2 infections were conducted in a biosafety level 3 facility.

To determine viral titers, virus-containing cell media were serially diluted (10^−2^ to 10^−5^) with DMEM into 96-well plates, 100 μL of each dilution was added in duplicate to 293T Ace2 cells (American Type Culture Collection) at 1 × 10^5^ cells per well in 24-well plates, and samples were incubated at 37 °C in 5% CO_2_ for 1 h, with rocking every 15 min to prevent cells from drying out. In parallel, plaquing media (MEM containing 2% FBS and 0.75% methylcellulose) was maintained at 37 °C in an incubator to decrease viscosity of the solution. After 1 h incubation, the virus-containing media were removed from the cells in the 24-well plates and 1 mL of plaquing media was added to each well. Plates were incubated at 37 °C in 5% CO_2_ for 3 days to allow plaque formation. On day 3, methylcellulose overlays were gently removed and cells were fixed by adding 1 mL 10% formaldehyde in PBS to each well. After incubation at room temperature for 20 min, the fixative was removed, plates were washed with dH_2_O, and 1 mL of 1% (w/v) crystal violet in 20% ethanol was added to each well. The crystal violet solution was removed after 15 min and plates were washed with dH_2_O until the plaques were visible. Plaques were only counted in wells containing 5–30 plaques. Titers were calculated in PFU/mL using the following formula: Titer (PFU/mL) = (number of plaques counted)/(dilution factor × volume plated).

To quantify viral genomic RNA in cell lysates, total RNA from mock or infected 293T Ace2 cells was isolated using the RNA NucleoSpin Kit (Machery-Nagel) and reverse-transcribed with random primers (Invitrogen) and the Improm-II reverse transcriptase system (Promega) at 42 °C for 1.5 hours. The resulting cDNAs were mixed with the SARS-CoV-2 spike gene-specific primers or β-actin primers (Table S1) (Integrated DNA Technologies) along with Perfecta SYBR Green SuperMix Low 6-Carboxy-X-Rhodamine (Quanta Biosciences) and then amplified for 40 cycles (each cycle 30 s at 94 °C, 40 s at 55 °C, and 20 s at 68 °C) in a Bio-Rad CFX96 Touch Real-Time PCR Detection System (Hercules, CA) as described previously.^45^ The ΔCT values were calculated using β-actin mRNA as the internal control. The ΔΔCT values were determined using control samples as the reference value. Relative levels of mRNAs were calculated using the formula 2(^−ΔΔCT^).^55^

### Cytotoxicity assays

293T Ace2 cells grown in opaque-walled 96-well plates were treated with DMSO or compounds at the desired concentration for 72 hours. After removing culture media, 100 µL of CellTiter-Glo Reagent (Promega) and 100 µL of PBS were added to each well, then the plate was placed on an orbital shaker for 2 minutes to induce cell lysis. The plate was incubated at room temperature for an additional 10 minutes before recording luminescence in a plate reader (Biotek Synergy HTX) at an integration time of 1 second per well.

### Microscale thermophoresis of ligand binding

RNA containing the SARS-CoV-2 pseudoknot (Table S1) was labelled with Cy5 at the 3′-end for microscale thermophoresis (MST).^56^ RNA was incubated at room temperature in 100 mM sodium acetate buffer (pH 5.0) with 100 mM NaIO_4_ to oxidize the 3′-terminal ribose, quenched with KCl, and purified by ethanol precipitation before incubation with Cy5 monohydrazide (Sigma, Cytiva PA15121) in 100 mM sodium acetate buffer (pH 5.0) for 4 hrs at room temperature with gentle agitation. Dye-labelled RNA (labelling efficiency ∼90%) was purified by gel purification and folded at ∼500 nM in MST measurement buffer (50 mM HEPES, 100 mM KCl, and 10 mM MgCl_2_, and 0.05% Tween).. MST measurements (Monolith NT.115, NanoTemper Technologies) were done with 150 nM RNA and 5% DMSO, with varying ligand concentrations. Fluorescence intensity was not affected by the ligand. Concentration-dependent curves (Fig. S3) were fit to a 1:1 binding model^57^ using MO.Affinity Analysis (v2.2.3) and Igor Pro (9.0.5.1) to determine *K*_D_; error in *K*_D_ was estimated by boot-strap analysis.

## Acknowledgements

We thank the Digital Alliance of Canada for providing access to computational resources for this project. We thank Dr. Darryl Falzarano (Vaccine and Infectious Disease Organization, University of Saskatchewan) for providing the SARS-CoV-2 isolate used in this work. We thank Dr. Darren Derksen (Chemistry, University of Calgary) for helpful discussions. We thank Dr. Marek Michalak lab (Biochemistry, University of Alberta) for access to the MST instrument.

## Author contributions

Conceptualization: MTW, JAT, and TCH; Methodology: RVS, SIO, and SA; Resources: KN, SMI, SM, and ER; Investigation: KN, RVS, SMI, ER, SM, SIO, SA, and JAP; Formal Analysis: KN, RVS, SMI, and SM; Visualization: KN, RVS, and MTW; Writing—original draft: KN, RVS, TCH, and MTW; Writing—review & editing: all authors; Supervision: MTW, JAT, and TCH; Project Administration: MTW; Funding Acquisition: MTW, JAT, and TCH.

## Supporting Information

**Table S1:**
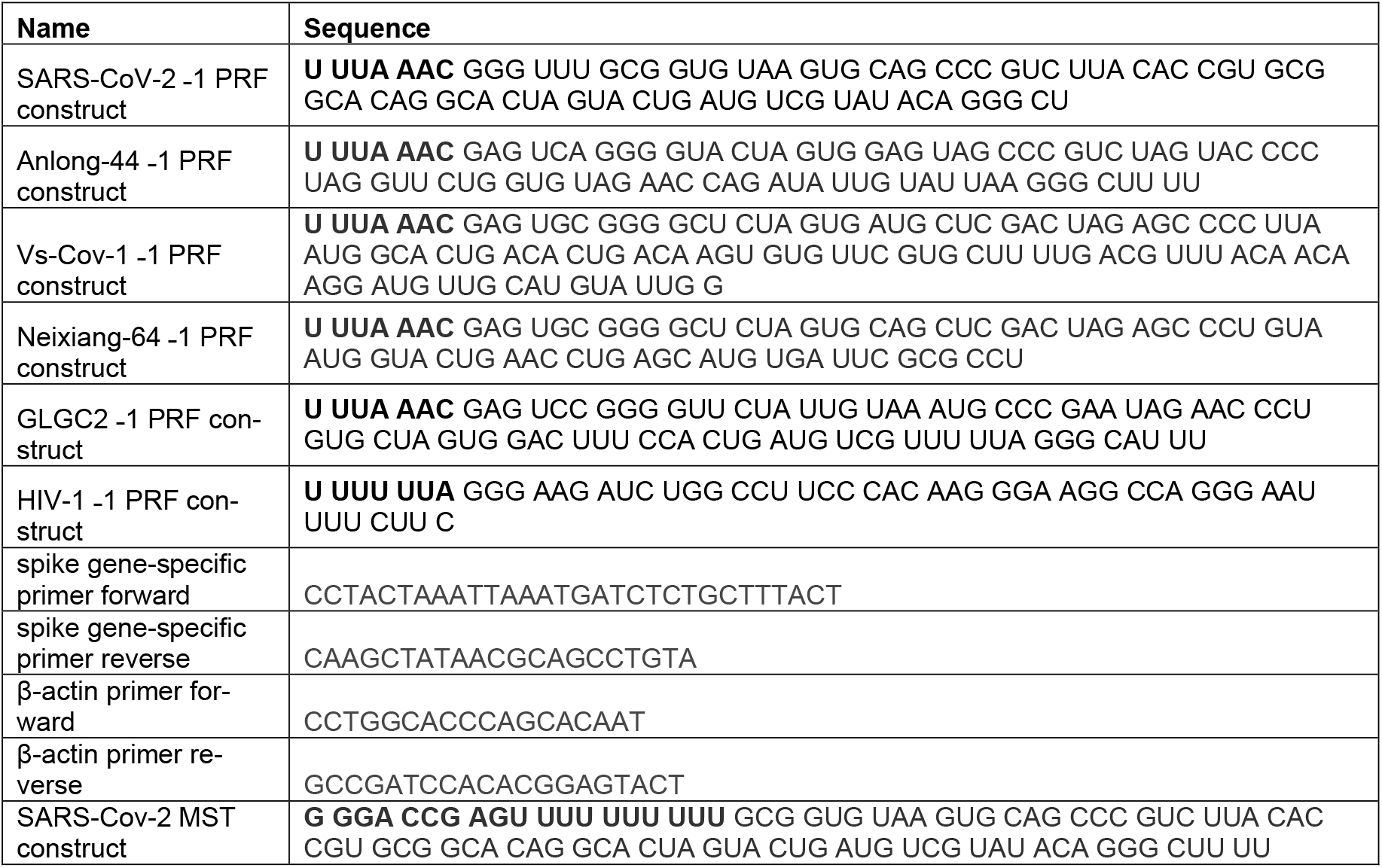
RNA sequences for −1 PRF constructs and DNA primer sequences.

**Fig. S1:**
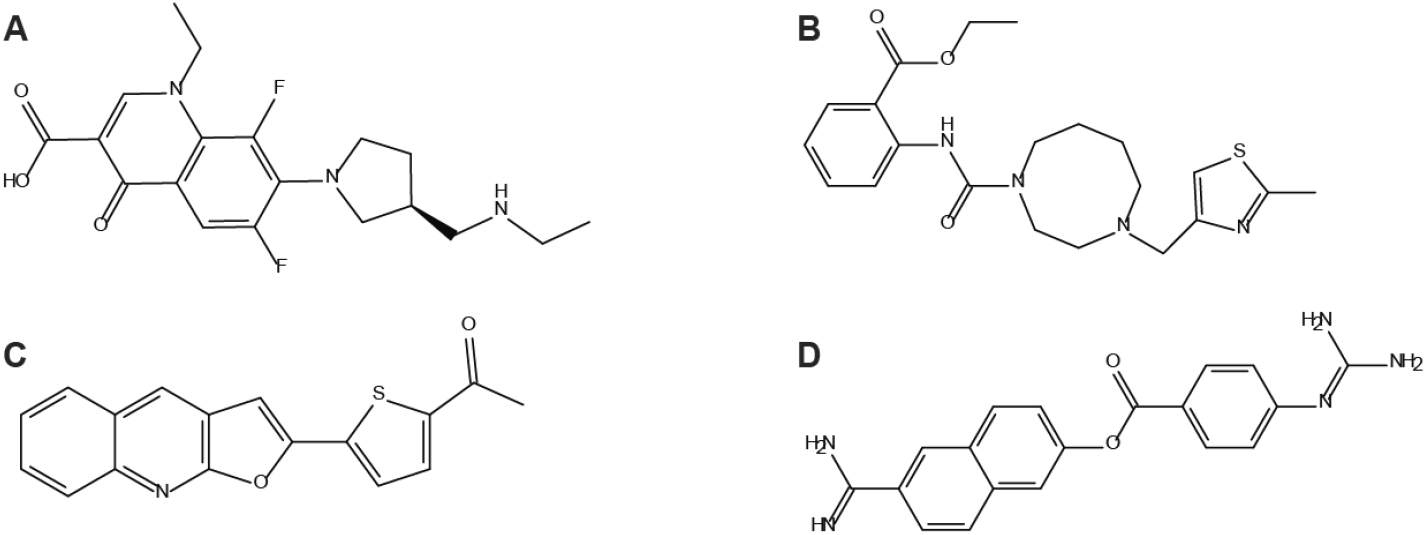
Structures of anti-CoV −1 PRF inhibitors reported previously. (A) Merafloxacin (ref. 14), (B) MTDB (ref. 19, 34), (C) KCB261770 (ref. 31), (D) nafamostat (ref. 17).

**Fig. S2:**
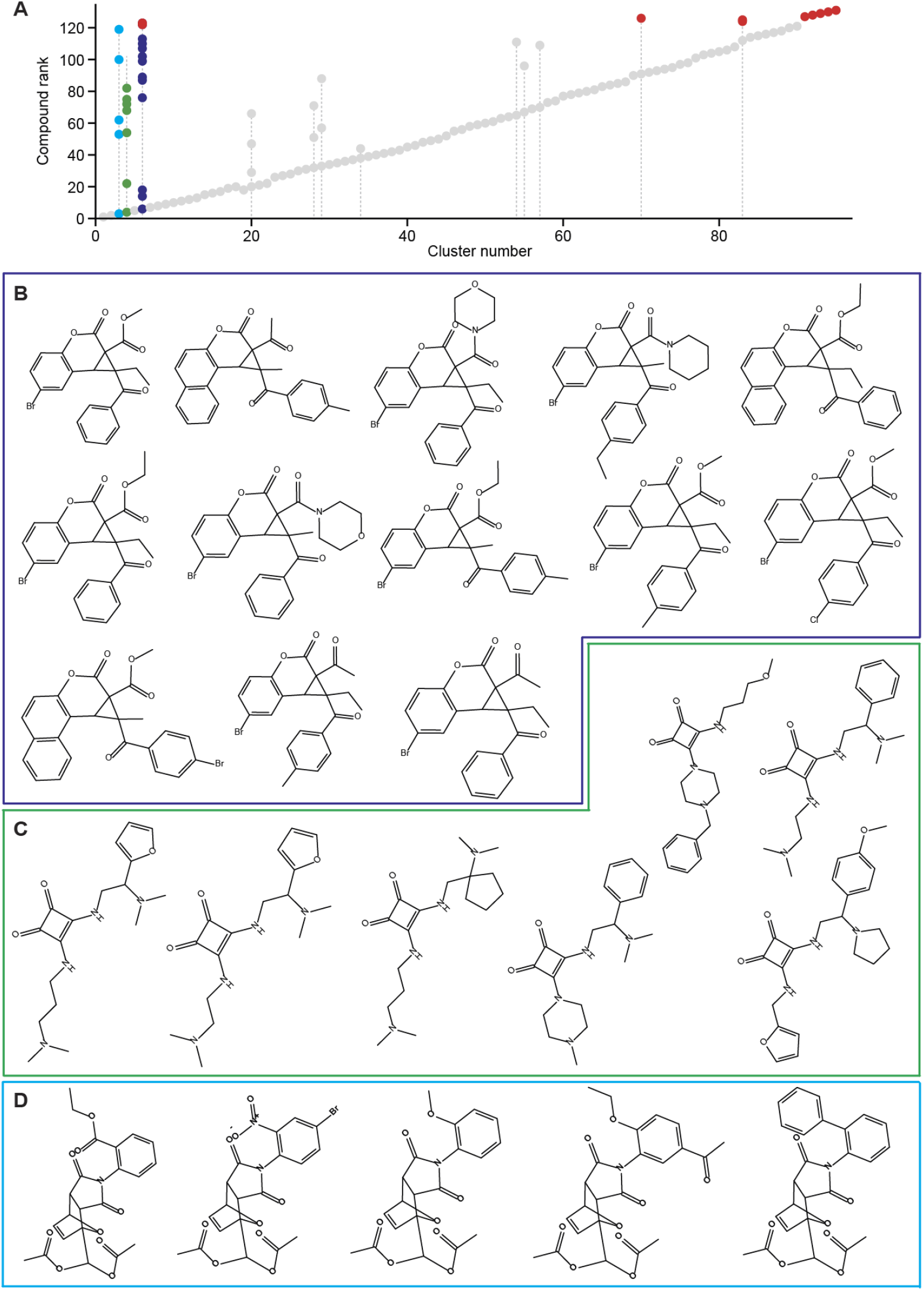
Chemical fingerprint cluster analysis of inhibitors. (A) The chemical fingerprints of all compounds tested in Fig. 3A were analyzed with Molecular Operating Environment (“graph 3-point pharmacophore” function applied to 2D molecular graphs) and clustered (using the “Tanimoto superset/subset” metric in MOE with similarity cutoff of 85% and overlap cutoff of 50%). The inhibitor rank is plotted as a function of cluster number, indicating that most compounds (grey) were chemically distinct (part of clusters with only 1 member), but some fingerprint clusters were selected multiple times by the docking algorithm. Top 10 compounds shown in red. (B–D) Structures of compounds from selected clusters with more than 1 member. (B) Compounds from cluster colored blue in panel A. (C) Compounds from cluster colored green in panel A. Compounds from cluster colored cyan in panel A.

**Fig. S3:**
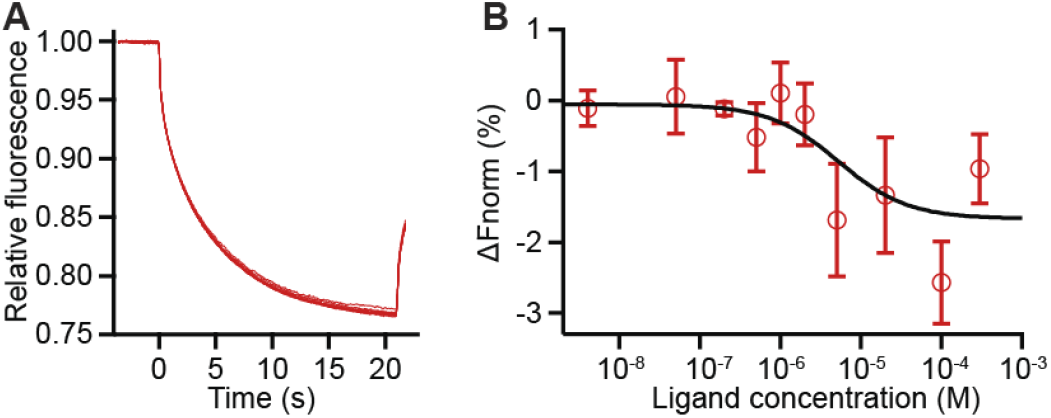
Binding of ligand 2 to SARS-Cov-2 frameshifting pseudoknot measured by MST. (A) Time-course of fluorescence during MST measurements for different concentrations of compound 2 (4 nM to 300 µM). (B) Concentration dependence of MST signal (red) fit to 1:1 binding model (black) yields an estimated *K*_D_ of 3 ± 2 μM, with relatively large uncertainty owing to low signal to noise ratio. Error bars represent s.e.m. from 3 independent measurements of different RNA samples.

